# Preclinical development of non-viral gene therapy for patients with advanced pancreatic cancer

**DOI:** 10.1101/2022.12.13.520242

**Authors:** Odile Barbey, Hubert Lulka, Naima Hanoun, Hafid Belhadj-Tahar, Fabienne Vernejoul, Gilles Cambois, Michèle Tiraby, Louis Buscail, Fabian Gross, Pierre Cordelier

## Abstract

Pancreatic ductal adenocarcinoma remains one of the greatest challenges in oncology for which therapeutic intervention is urgently needed. We demonstrated that the intratumoral gene transfer of somatostatin receptor 2, to combat tumor aggressiveness, or of deoxycytidine kinase and uridylate monophosphate kinase, to sensitize to gemcitabine chemotherapy, has antitumoral potential. Here, we describe the development of CYL-02 non-viral gene-therapy product, that comprises a DNA-plasmid encoding for the three aforementioned genes complexed with PolyEthylEnimine (22 kDa). In this work, we performed preclinical toxicology, biodistribution and therapeutic activity studies of CYL-02 in experimental models of pancreatic cancer. We demonstrated the safety of CYL-02 and defined the maximal tolerated dose in two animal species. CYL-02 co-administrated with gemcitabine did not increase gemcitabine toxicity. Biodistribution studies revealed that CYL-02 is rapidly cleared from blood following intravenous administration, and sequestered in tumors following intratumoral injection. CYL-02 drives the expression of therapeutic genes in cancer cells and strongly sensitizes tumor cells to gemcitabine, with significant inhibition of tumor cells dissemination. This study was instrumental for the later use of CYL-02 in patients with advanced pancreatic cancer, demonstrating that rigorous and thorough preclinical investigations are informative for the clinical transfer of gene therapy against pancreatic cancer.

**GRAPHICAL ABSTRACT:** 

## INTRODUCTION

Pancreatic ductal adenocarcinoma (PDAC) will soon become the second cause of death by cancer worldwide ^1^. Due to a long silent clinical phase during tumor development and to the absence of early markers that delay diagnosis, the prognosis of this cancer is very poor. When surgery is not possible, therapeutic options are few and ineffective^2^. Patients are most often treated with Folfirinox or gemcitabine chemotherapy that ameliorates few clinical parameters including pain intensity, analgesic consumption and weight loss. Tumor volume is nearly unchanged and survival remains extremely short. Clearly, more effective treatments are urgently needed to alleviate the dismal prognosis of PDAC. Thus, the moderate activity of standard therapies strongly encourages new translational research programs such as gene therapy to potentiate the antitumoral effect of chemotherapy.

With this in mind, we demonstrated during the last two decades the antitumoral potential of the somatostatin receptor subtype 2 (SSTR2) in PDAC experimental models. SSTR2 expression is lost in 95% of PDAC tumors^3^, and restoring SSTR2 using non-viral gene transfer results in a strong bystander antitumoral effect that is antiproliferative, pro-apoptotic, anti-angiogenic, and anti-metastatic^4–9^. In addition, SSTR2 was found by others to sensitize tumor cells to gemcitabine^10^. In parallel, we explored PDAC tumors cells to identify candidate that may help defeat resistance to gemcitabine. DeoxyCytidine Kinase (DCK) phosphorylates gemcitabine to gemcitabine diphosphate in a rate-limiting step. Loss of expression of this enzyme was recently associated with acquired resistance to gemcitabine in pancreatic cancer cells, in preclinical models ^11^, and in patients ^12^. We demonstrated that delivering *DCK::UMK* fusion gene, encoding for both DCK and Uridylate Monophosphate Kinase (UMK, also named NMPK for nucleoside monophosphate kinase), using non-viral vectors overcomes PDAC-derived cells resistance to gemcitabine ^13^. Thus, as opposed to many trials for PDAC treatment in which new agents are combined with gemcitabine simply because it is a standard of care, there is a strong rationale to deliver *SSTR2, DCK* and *UMK* coding sequences and to treat advanced pancreatic cancer tumors with gemcitabine as chemotherapy. In this work, we describe the preclinical development of the CYL-02 gene therapy product, encoding for *SSTR2, DCK* and *UMK*, that was later used in combination with gemcitabine. in phase 1^14^ and phase 2^1^ clinical trials in patients with advanced PDAC.

## RESULTS

### Preclinical assessment of the toxicity of the CYL-02 gene therapy product

For this study, we generated plasmid DNA encoding for *SSTR2, DCK* and *UMK* coding sequence which expression is targeted to tumor cells as described in Materials and Methods. Therapeutic DNA is complexed with 22 KDa polycationic PolyEthylEnimine (in vivo-jetPEI®) to give CYL-02 non-viral gene therapy product (GTP) as described before^14^., We generated 2 pre-GMP and 1 GMP batches as part of the pharmaceutical development of CYL-02. We first aimed to determine the largest amount of CYL-02 at which no detectable adverse effects occur in animal models (No Observable Adverse Effect Level, NOAEL), starting with 7500 μg/kg as the toxic dose of PEI in mice^15^. Thus, female and male C57Bl/6 mice were injected intravenously (i.v.) with 250μg/kg, 1500μg/kg and a maximal dose of 7500μg/kg of the GTP in pre-GMP grade. Control mice received CYL-02 excipient (5% glucose) as placebo as this is the resuspension solution of CYL-02, and mice were monitored for 5 weeks (Supplemental figure 1A). No mortality, nor weight loss or changes in Aspartate-Amino-Transferase (ASAT), ALAT (Alanine aminotransferase), lactate dehydrogenase (LDH) or creatinine levels were found in mice receiving an i.v. injection of 250μg/Kg or 1500μg/kg of the GTP (Supplemental Figure 2A-I). On the other hand, half of the mice, two male mice (40%) and 3 female mice (60%), died within 24 hours following i.v. injection of the highest dose of CYL-02. We performed pathology analysis to better characterized the toxic effect following injection of CYL-02 in mice. Three out of 15 (20%), and 4 out of 30 (13%) male and female mice show mononuclear cells aggregates in the liver, respectively, regardless of the dose of CYL-02 (Figure 1A). Alveolar atelectasis was identified in the lungs of 11/30 (37%) and of 5/30 (17%) of male and female mice, respectively, regardless of the dose of CYL-02 (Figure 1B). Last, one mice that received the highest dose of CYL-02 showed evidence of kidney congestion and small intestine autolysis (Figures 1C and 1D), as a result of agony before mouse euthanasia. Taken together, i.v. injection of CYL-02 is safe in mice with a maximal tolerated dose (MTD) of 1500μg/kg.

**Figure 1.**
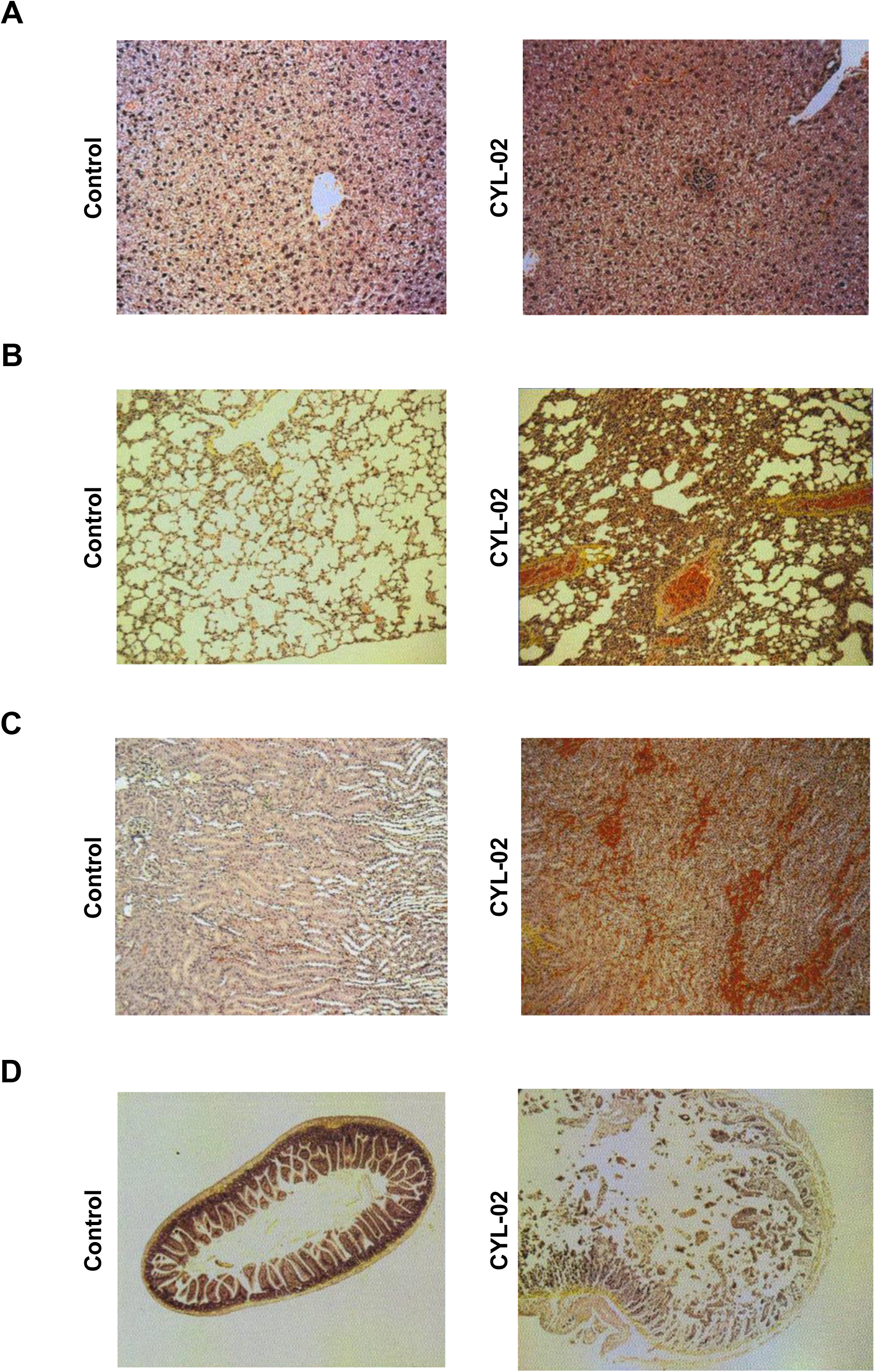
Toxicology study following a single i.v. injection of CYL-02 in mice. 250μg/kg, 1500μg/kg and 7500μg/kg of first generation CYL-02 was administered i.v. in a total of 45 C57Bl/6 mice from both sex. Five weeks later, mice were enthanased and analyzed for toxic events associated with gene therapy administration in liver (**A**), lung (**B**), kidney (**C**) and bowel (**D**).

We then extended the toxicity study to a non-rodent animal model following the guidelines of the International Council for Harmonisation (ICH) of technical Requirements for Pharmaceuticals for Human Use (ICH M3; S6 and S9). Here, pre-GMP grade CYL-02 was administrated by the intended route of administration in humans, *i.e*. in tumors. We generated orthotopic PDAC tumors in lagomorphs (Syrian gold hamsters) as described in Materials and Methods and Supplemental figure 1B. Seven days later, exponentially growing tumors were injected with a starting dose of 500μg/kg of pre-GMP grade first generation of CYL-02, that was decided based on the MTD identified following i.v. injection of the GTP in mice, after correction with a clearance factor for injection in hamster. Control animals received 5% glucose. Animals received 80mg/kg of gemcitabine by intraperitoneal (i.p.) route at days 2, 4 and 6 following intratumoral gene transfer. No animal died during the experiment, with only mild body weight loss in hamsters receiving gemcitabine, or CYL-02 and gemcitabine (−21%±9%, *p*<0.01, Figure 2A). We next evaluated the toxicity associated with repeated injections of CYL-02, as this may induce an unwanted and adverse immune response in patients. Thus, both male and female C57B/6 mice were injected i.v. with 250 μg/kg of CYL-02 at day 0, 15 and 25 (Supplemental figure 1A). All animals injected survived the experiment, and no organ-specific toxicity was detected. We next performed a complementary experiment by injecting 250 μg/kg of CYL-02 i.v. in the same murine model, followed by subcutaneous injection of 250 μg/kg of the gene therapy product at day 15, 25 and 40. Here again, animals were examined for 48h after each injection for signs of systemic and cutaneous toxicity, and no local reactions were observed after iterative subcutaneous injection preceded or not intravenous injection.

**Figure 2.**
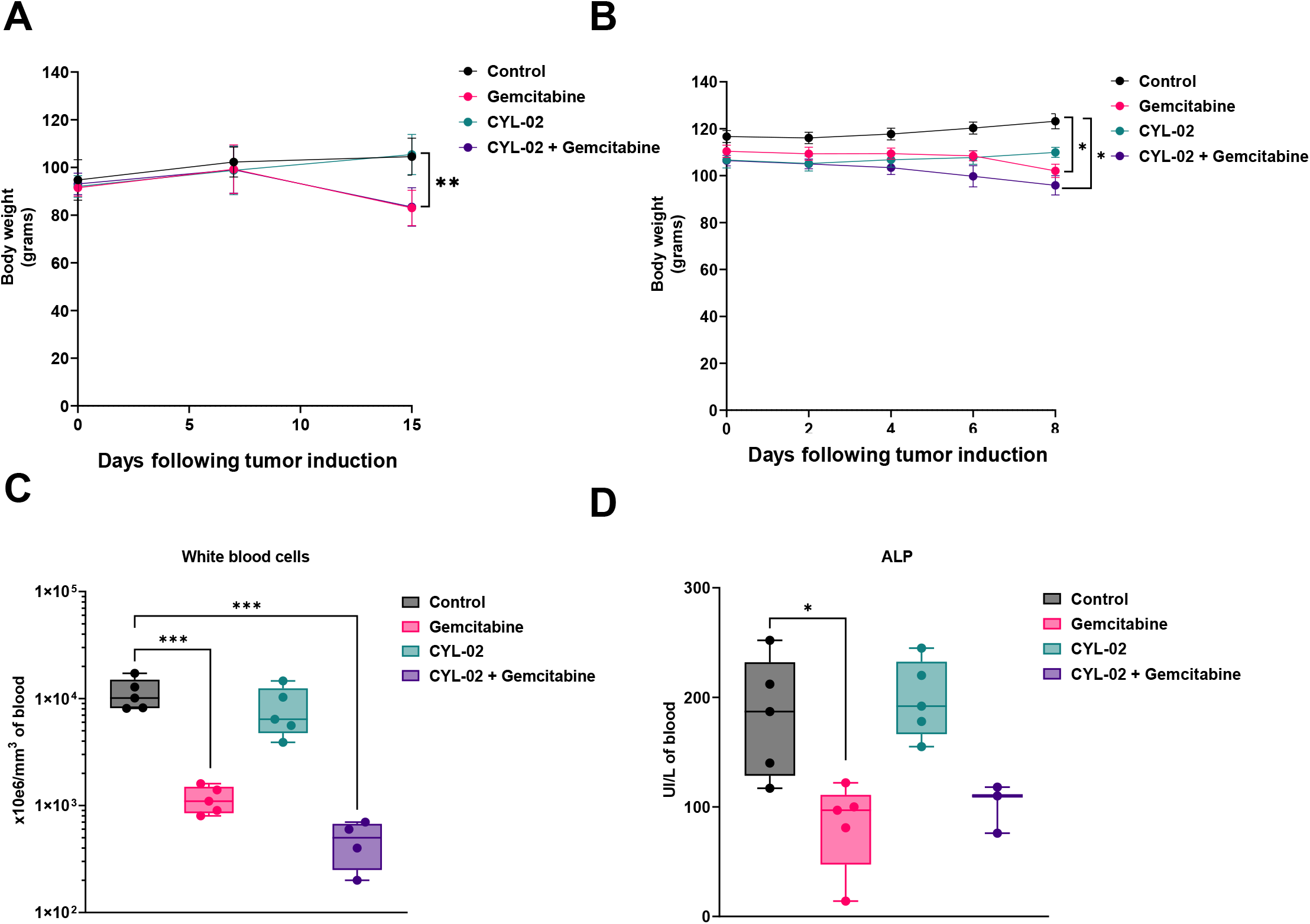
Toxicology study following a single intratumoral injection of CYL-02 in orthotopic pancreatic tumors in Syrian gold hamsters. Experimental orthotopic PDAC tumors were induced as described in Materials and Methods. Eight days later, 500μg/kg of first (**A**) or 900μg/kg of second (**B**) generation CYL-02 was administered in exponentially growing tumors, when control animals received 5% glucose. Gemcitabine was given at 80mg/kg i.p. every 2 days for a week. NaCl9^0/00^ was given i.p. as control. Hamster body weight was monitored from tumor induction, up to 7 days following treatment. n=5 hamsters were used per group. White blood cells (**C**) and alkaline phosphatase (**D**) monitoring in hamsters receiving an intratumoral injection of CTL-02 combined with gemcitabine treatment. *: p<0.05, **: p<0.01, ***: p<0.005.

In this first version of the GTP, the resistance gene for bacterial selection (*NEO*) was co-expressed with *SSTR2* cDNA due to the presence of an internal ribosomal entry site (IRES). To minimize the risk associated with the expression of a resistance gene expression in recipient and finally in humans, we generated a second version of CYL-02 in which the *NEO* gene expression is driven by a bacterial promoter. We next performed “bridging” toxicology studies to address whether first and second generation of CYL-02 behaved the same in hamsters. Thus, PDAC tumors generated in hamsters were injected with 900μg/kg of pre-GMP grade CYL-02 of second generation. This dose corresponds for hamsters to the MTD identified in mice. Blood and urine were sampled for three animals per group15min, 30min, 1h, 3h, 6h, 12h, 18h and 24h after injection. We did not observe any evidence of acute nor systemic toxicity related to CYL-02 intratumoral injection in experimental tumors in hamsters.

As for first generation CYL-02, we next injected second generation, GMP-grade CYL-02 in exponentially growing PDAC tumors in hamsters in combination with gemcitabine to fully capture the toxicity related to one injection of GTP combined to chemotherapy administration. GMP-grade, second generation CYL-02 administration was not associated with animal death, with the exception of one hamster that also received gemcitabine (1/5, 20%). This death was preceded by diarrhea and the alteration of the general condition resulting in a significant weight loss (−15%) and lethargy. Autopsy revealed significant tumor invasion in the spleen, stomach, liver, bowel and colon by regional extension. As shown in Figure 2B, animals receiving either CYL-02 or placebo gained weight with a similar trend (+9%±2.8% vs+8.9%±6.2%, respectively). On the other hand, significant weight loss was measured when gemcitabine was administered alone (−4.1±2%, *p*<0.05) or in combination with CYL-02 (−3.6%±1%, *p*<0.05). We identified leukopenia in the gemcitabine group and in combination with CYL-02, as white blood cells count significantly dropped as compared to control (Figure 2C, −89%±2%, *p*<0.005 and −94%±3%, *p*<0.005, respectively). To a lesser extent, we also measured a significant decrease in alkaline phosphatase (ALP) in hamsters treated by gemcitabine only (Figure 4D, −56±11%, *p*<0.05). No changes in red blood cells and platelets count, and ASAT, ALAT and creatinine level were identified between the different groups (Supplemental Figure 3A-D). Pathology examination of the organs at autopsy did not revealed significant alterations in the brain, lung, heart, bladder and muscles of the animals between the different study groups. However, we found systematic peritoneal alterations due to repeated laparotomies and i.p. injections. In addition, all study groups evidenced testicular inflammation, as a marker of abdominal tumor development. Last, one hamster out of five (20%) receiving gemcitabine alone demonstrated renal parenchyma alteration and central testicular necrosis without bladder involvement.

**Figure 3.**
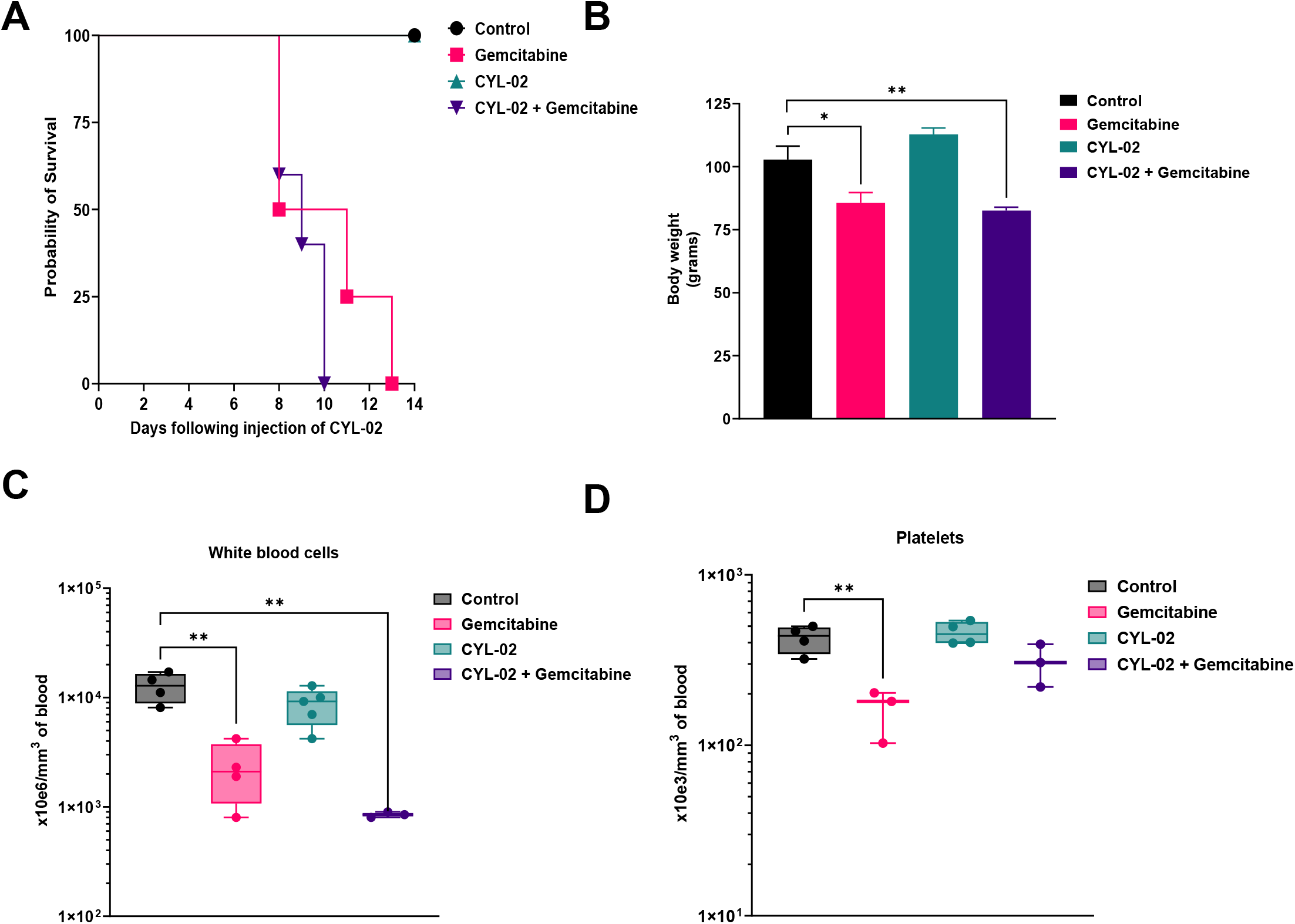
Toxicology study following two intratumoral injection of CYL-02 in orthotopic pancreatic tumors in Syrian gold hamsters. Experimental orthotopic PDAC tumors were induced as described in Materials and Methods. 500μg/kg of second generation CYL-02 was administered at days 0 and 7 in exponentially growing tumors, when control animals received 5% glucose. Gemcitabine was given at 80mg/kg i.p. every 2 days for a week following each injection. NaCl9^0/00^ was given i.p. as control. Hamster body weight was monitored up to 14 days following the first intratumoral injection of the gene therapy product. n=5 hamsters were used per group. **A**. Probability of survival between the different experimental groups. **B**. Hamster body weight, 15 days following the first intratumoral gene trransfer. White blood cells (**C**) and platelets (**D**) monitoring in hamsters receiving two cycles of intratumoral injection of CTL-02 combined with gemcitabine treatment. *: p<0.05, **: p<0.01.

**Figure 4.**
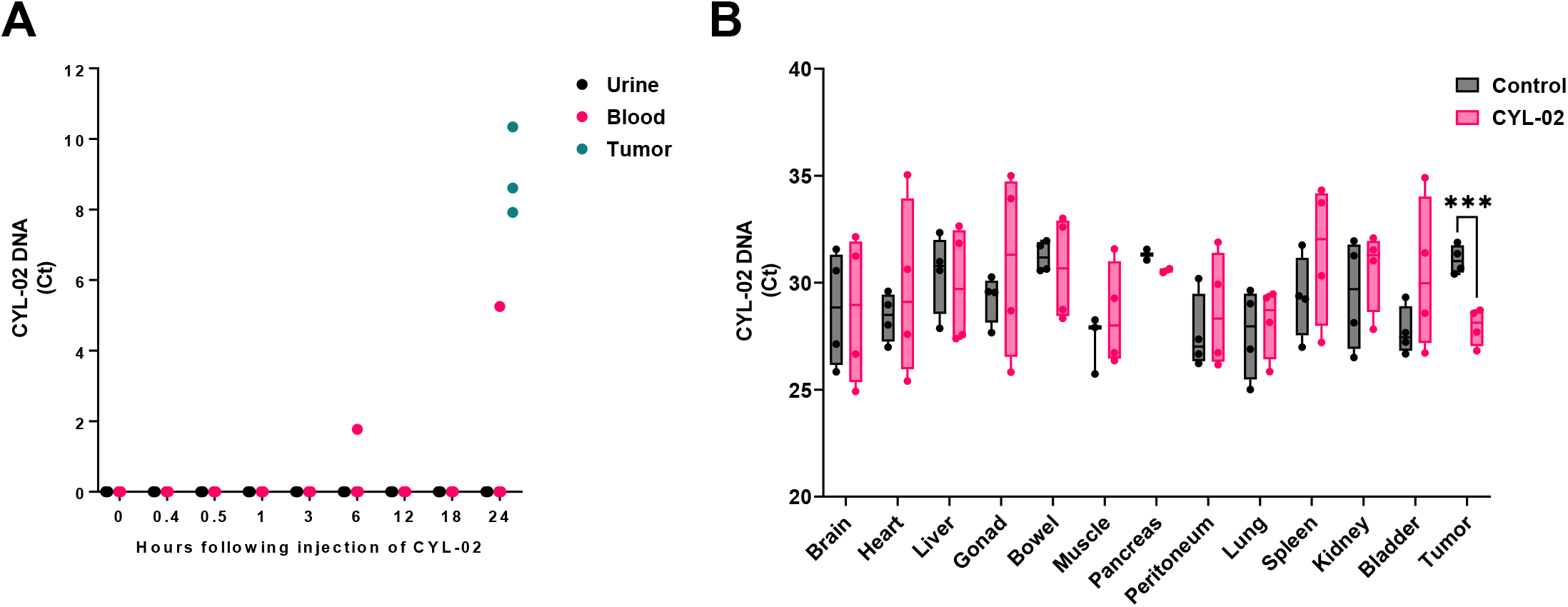
Biodistribution study following intratumoral injection of CYL-02 in orthotopic pancreatic tumors in Syrian gold hamsters. Experimental orthotopic PDAC tumors were induced as described in Materials and Methods. **A**. 900μg/kg of second generation CYL-02 was administered in exponentially growing tumors, and urine and blood were sampled at the indicated time after gene transfer. As control, tumors were sampled and analyzed 24 hours following intratumoral gene transfer of CYL-02. CYL-02 DNA was detected by qPCR for neomycin gene (not expressed in eukaryotic cells). Data are expressed as Ct of 3 biological replicates. **B**. 900μg/kg of second generation CYL-02 was administered at days 0 and 7 in exponentially growing tumors, when control animals received 5% glucose. Gemcitabine was given at 80mg/kg i.p. every 2 days for a week following each injection. NaCl9^0/00^ was given i.p. as control. CYL-02 DNA was detected in the indicated organs and in tumours by qPCR for neomycin gene (not expressed in eukaryotic cells). Data are expressed as Ct ± SD for Neomycin gene of 5 biological replicates per group with 3 experimental replicates. ***: p<0.005.

We then questioned the safety of repeated intratumoral injection of CYL-02 following the treatment scheme we wanted to test in humans in clinical trial. Thus, pre-GMP grade CYL-02 was administered into tumors at days 0 and 7, and gemcitabine was given i.p. at days 2, 4 and 6 days (first cycle) and 9, 11 and 13 days (second cycle) after the first injection of the GTP. While CYL-02 injection alone was safe, we found that animal survival was significantly shortened when receiving gemcitabine (*p*<0.01) or CYL-02 + gemcitabine (*p*<0.01) as compared to control (Figure 3A). Gemcitabine toxicity also translated into significant body weight loss when administered alone (−17%±10%, *p*<0.05) or in combination with CYL-02 (−20%±4%, *p*<0.01), as compared to control (Figure 3B). Here again, CYL-02 is safe as it did not alter animal body weight (Figure 3B). Gemcitabine and CYL-02 combined to gemcitabine treatment resulted in leukopenia (Figure 3C), and we also identified a drop in platelets count in hamsters receiving the chemotherapy only (Figure 3D). Numbers of red blood cells and level of ASAT, ALAT, LDH and creatinine remained unchanged between the different groups (Supplemental figure 4A-E). Thus, these data demonstrate the safety of second generation, GMP-grade CYL-02 in hamster following 2 intratumoral injections in experimental tumors when combined with gemcitabine. Collectively, toxicity studies revealed that the MTD for CYL-02 in mice is of 1500 μg/Kg corresponding to 900μg/kg for Syrian gold hamsters, when adjusted with a clearance factor. In both animal models, CYL-02 is safe, even following repeated administration, and does not aggravate gemcitabine toxicity.

### Preclinical study of the biodistribution of the CYL-02 gene therapy product

We performed a first set of studies to address the biodistribution of the first version of pre-GMP CYL-02 in female and male mice receiving i.v. 250μg/kg, 1500μg/kg or 7500μg/kg of the gene therapy product. Eleven organs were analyzed by quantitative PCR (qPCR) for CYL-02 DNA detection, up to 6 weeks following injection. Table 1 shows that CYL-02 DNA is readily detectable in all organs tested, 6-hours post-injection of female mice. Twenty-four hours later, we sampled blood and urine from animals receiving 250μg/kg of the gene therapy product and found that only one fourth of the urine tested were positive for CYL-02, when all blood samples tested were negative (data not shown). After 7 days, CYL-02 was detected inconsistently in the lung, liver, kidney and the heart of animals that received 250 and 1500 μg/kg of the GTP. In addition, female mice receiving the highest dose of CYL-02 (7500μg/kg) showed positivity in muscles and brain. By 2 weeks, most male and female mice were free of CYL-02 (Tables 1 and 2), regardless of the dose injected. Collectively, these data show that CYL-02 is only transiently detectable for 28 days in lung, heart, liver and pancreas and disappears 35 days following i.v. injection in mice, respectively.

**Table 1:** Analysis of the biodistribution of first generation CYL-02 in female mice using qPCR following i.v. injection at the indicated dose. -: negative, +: 2.5<ddCt<5, ++: 5<ddCt<10, +++: 10<ddCt<20, ++++: ddCt>20. ddCt=[(Ctneo-CtGAPDH) in organ from the injected group] - [(Ctneo-CtGAPDH) in organ from the non-injected group). H: hours, D: day.

| 250µg/kg | Lung | Kidney | Bowel | Heart | Muscle | Brain | Peritoneum | Pancreas | Spleen | Liver | Bladder | Gonads |
| --- | --- | --- | --- | --- | --- | --- | --- | --- | --- | --- | --- | --- |
| D7 | - | - | - | - | - |  | - | - | - | - | - | - |
| D14 | - | - | - | - | - | - | - | - | - | - | - | - |
| D21 | - | - | - | - | - | - | - | - | - | - | - | + |
| D28 | - | - | - | - | - | - | - | + | - | + | - | - |
| D35 | - | - | - | + | - | + | - | - | - | - | - | - |
| **1500µg/Kg** |  |  |  |  |  |  |  |  |  |  |  |  |
| D7 | ++ | ++ | - | ++ | - |  | - | - | - | + | - | + |
| D14 | + | - | - | - | - | - | - | - | - | + | - | + |
| D21 | - | - | - | + | - | - | - | - | - | + | - | + |
| D28 | + | - | - | + | - | - | - | - | - | - | - | - |
| D35 | - | - | - | - | - | - | - | - | - | - | - | - |
| **7500µg/Kg** |  |  |  |  |  |  |  |  |  |  |  |  |
| H6 | ++++ | +++ | ++ | +++ | +++ | +++ | +++ | +++ | +++ | +++ | + | +++ |
| D7 | ++ | - | - | + | + | ++ | - | - | - | - | - | - |
| D14 | - | - | - | - | - | - | - | - | - | - | - | - |

**Table 2:**
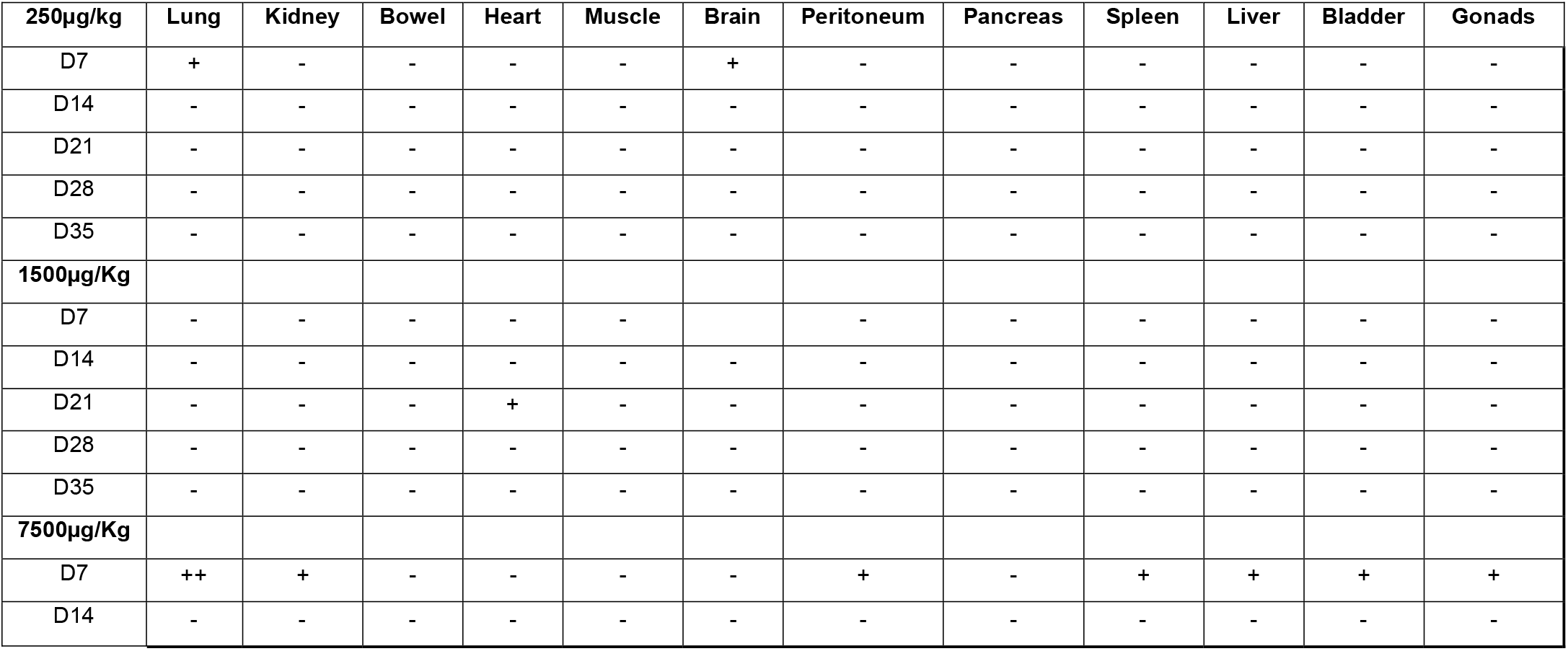
Analysis of the biodistribution of first generation CYL-02 in male mice using qPCR following i.v. injection at the indicated dose. -: negative, +: 2.5<ddCt<5, ++: 5<ddCt<10, +++: 10<ddCt<20, ++++: ddCt>20. ddCt=[(Ctneo-CtGAPDH) in organ from the injected group] - [(Ctneo-CtGAPDH) in organ from the non-injected group). D: day.

We further analyzed the biodistribution of the first version of the GTP in hamsters following intratumoral injection of 500μg/kg of pre-GMP CYL-02 (Day 0), followed by treatment with gemcitabine *i.p*. at days 2, 4 and 6. Organs (spleen, liver, lung, kidney and bowel) and tumors were sampled 7 days following gene transfer. Table 3 shows that CYL-02 levels are high in tumors, and in the liver and, to a lesser extent, the spleen of animals receiving CYL-02 and gemcitabine. We then moved to second generation CYL-02 as mentioned before. Hamster PDAC tumors were injected with 900μg/kg of pre-GMP CYL-02 and urine and blood were sampled 15min, 30min, 1h, 3h, 6h, 12h, 18h and 24h later. Tumors were sampled 24 hours following intratumoral injection of the GTP. Figure 4A shows that CYL-02 is not detected in urine (0/24), and rarely in the blood (2/6); indeed, CYL-02 was found in the blood of 1/3 hamster 6 hours (corresponding to 0.1% of the dose injected) or 24 hours (corresponding to 2.3% of the dose injected) following intratumoral injection, when tumors showed high levels of the gene therapy product. We conclude that CYL-02 is largely sequestered in the tumor with an inconstant and delayed presence of a small quantity of the vector in the vascular compartment. We then performed repeated intratumoral injection of second generation, pre GMP CYL-02 in exponentially growing hamsters PDAC tumors followed by i.p. injection of gemcitabine. Fourteen following the first injection, hamsters were euthanized and CYL-02 expression was analyzed by qPCR in the tumor, pancreas, lung, liver, bowel, gonad, kidney, bladder, spleen, heart, striated muscle, brain and peritoneum. As shown in Figure 4B, significant increase in CYL-02 was identified in tumors from animal receiving the GTP, up to 2 weeks following intratumoral injection (+8.75±1.3-fold increase, *p*<0.001). Thus, we demonstrate that CYL-02 is cleared from organs 1 month following i.v. injection, and is sequestered in tumors up to 7 days following intratumoral injection with minimal, early, exposure in the blood.

**Table 3:** Analysis of the biodistribution of first generation CYL-02 in hamsters using qPCR, 7 days following intratumoral injection of the GTP. -: negative, +: 2.5<ddCt<5, ++: 5<ddCt<10, +++: 10<ddCt<20, ++++: ddCt>20. ddCt=[(Ctneo-CtGAPDH) in organ from the injected group] - [(Ctneo-CtGAPDH) in organ from the non-injected group).

|  | Lung | Kidney | Bowel | Spleen | Liver | Tumor |
| --- | --- | --- | --- | --- | --- | --- |
| **CYL-02** | - | - | - | + | ++ | ++ |
| **CYL-02 + gemcitabine** | - | + | - | ++ | +++ | +++ |

### Characterization of CYL-02 activity in preclinical models of PDAC

We next analyzed *SSTR2, DCK* and *UMK* gene expression following *in vitro* transfection of hamster pancreatic cancer cells. Thus, PC.1-0 cells were transfected with 1μg of second generation CYL-02 (that will now be designed as CYL-02). Control cells received 5% glucose. Forty-height hours later, therapeutic gene expression was quantified by reverse transcription followed by qPCR (RTqPCR), as described before^14^. Figure 5A demonstrates that CYL-02 strongly increases *SSTR2* (844-fold±307, *p*<0.01), *DCK* (249-fold±123, *p*<0.01) and *UMK* expression (516-fold±406, *p*<0.05) in transfected cells as compared to control cells.

**Figure 5:**
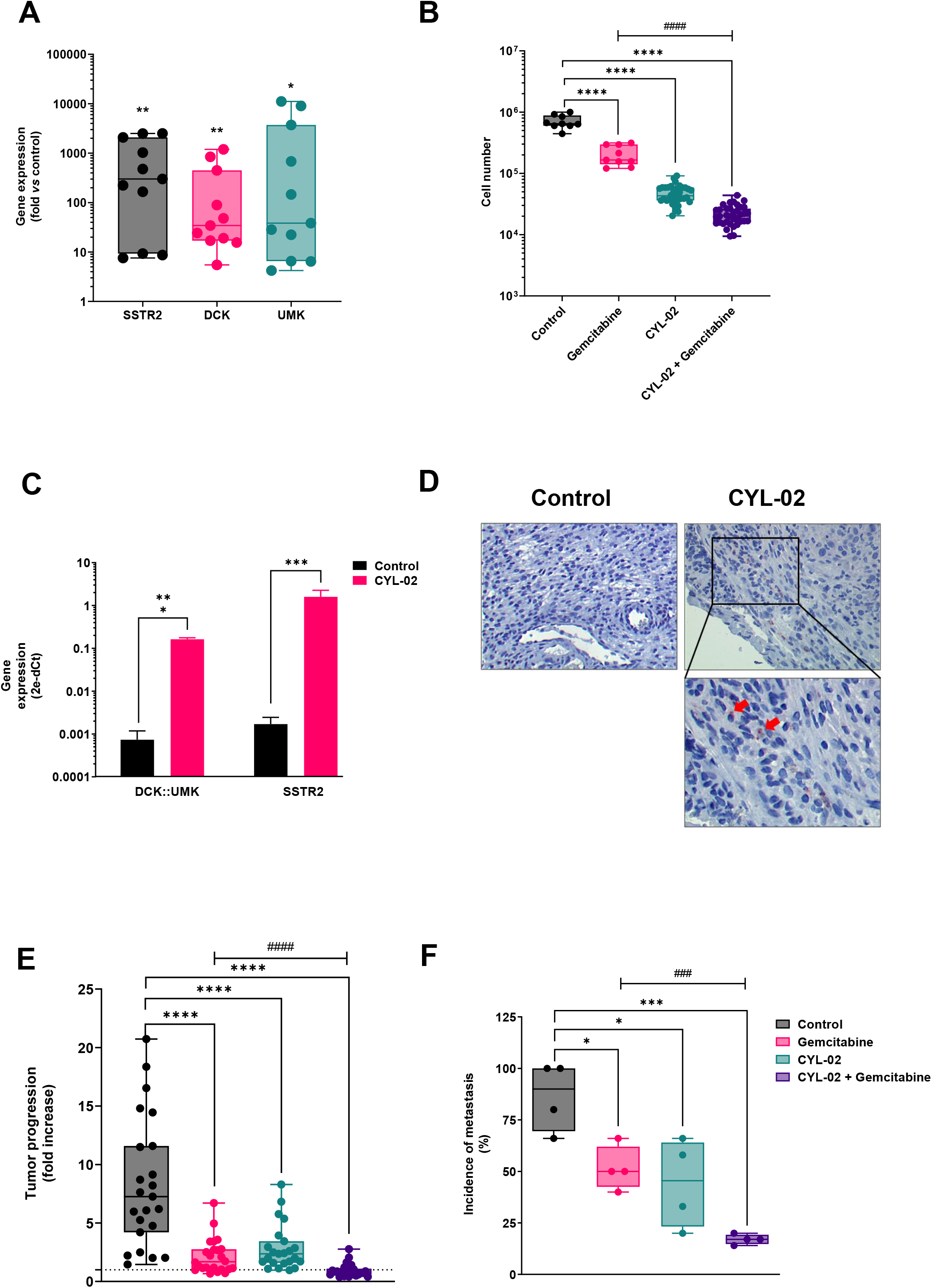
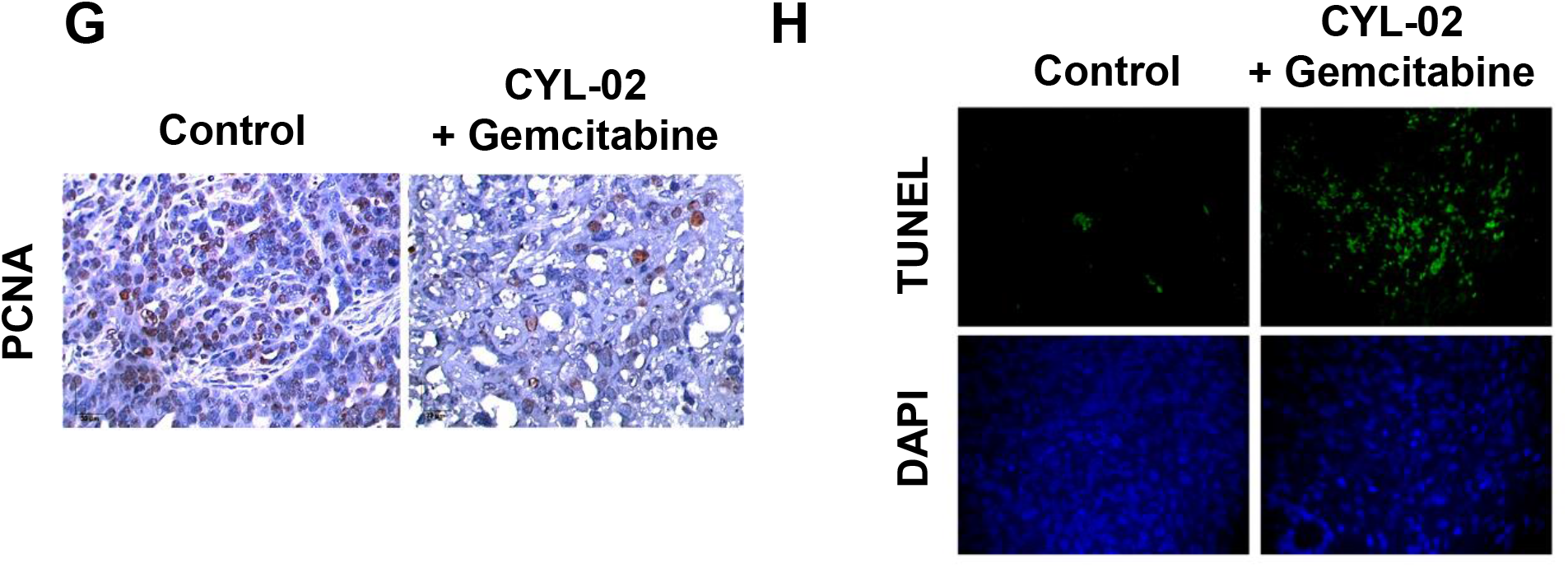
preclinical characterization of CYL-02 activity and antitumoral efficacy in orthotopic pancreatic tumors in Syrian gold hamsters. **A.** PC. 1-0 hamster pancreatic cancer cells were transfected with CYL-02 as described in Materials and Methods, and *SSTR2, DCK* and *UMK* gene expression analysis was performed as described elsewhere^14^. Results are mean of n=11 independent experiments performed in duplicate and expressed as mean fold change ± SD between control and transfected cells, using 18S as an internal control. **B**. PC.1-0 hamster pancreatic cancer cells were transfected with CYL-02 and treated by gemcitabine as described in Materials and Methods. Three days later, cells were counted. Results are mean ± SD. of 9 (control, gemcitabine) or 36 (CYL-02, CYL-02 + gemcitabine) independent experiments. C. Experimental orthotopic PDAC tumors were induced as described in Materials and Methods. Eight days later, 900μg/kg of second generation CYL-02 was administered in exponentially growing tumors, when control animals received 5% glucose. Gemcitabine was given at 80mg/kg i.p. every 2 days for a week. NaCl9^0/00^ was given i.p. as control. At the end of the experiments, mice were enthanased and (**C**) *DCK::UMK* and *SSTR2* gene expression was measured in control tumors and CYL-02-treated tumors, 8 days following gene therapy. Data are means ± SD of 5 biological replicates per group with 3 experimental replicates and expressed as arbitrary units (2^-ΔCt^ with ΔCt= CT(*DCK::UMK or SSTR2*) – CT(*18S*)). **D**. SSTR2 protein was detected by immunochemistry in control and CYL-02-treated experimental tumors 8 days following gene therapy. Fifteen fields were analyzed per condition. Data are representative of 5 biological replicates per group with 3 experimental replicates. All images: x40 original magnification. Scale bar is 30μm and valid for all images. Red arrows indicate cells positive for SSTR2. At the end of the experiment, tumor progression (**E**) and dissemination (**F**) were analyzed between the different groups. Results are mean fold ± ND of tumor progression in n=20 animals per group, sum of 8 independent experiments, and of tumor dissemination (n=20 for each condition, sum of 4 independent experiments). Dash line indicates no progression (fold=1). Following autopsy, tumors were sampled and analyzed for PCNA expression (**G**) and TUNEL assay (**H**). Fifteen fields were analyzed per condition. Data are representative of 5 biological replicates per group with 3 experimental replicates. All images: x40 original magnification. Scale bar is 30μm and valid for all images.

We then performed cell proliferation studies to document the activity of CYL-02. Thus, PC.1-0 cells were transfected with 1μg CYL-02, and treated or not by gemcitabine 48 hours later. Control cells received either 5% glucose (CYL-02 control) and/or NaCl 9^0/00^ (Gemcitabine control). Forty-eight hours later, cells were counted. Results shown in Figure 5B demonstrate that gemcitabine significantly inhibits the proliferation of PC.1-0 cells (−71%±12%, *p*<0.0001). Remarkably, CYL-02 transfection strongly inhibits hamster PDAC cells proliferation (−94%±5%, *p*<0.0001), and combination of CYL-02 and gemcitabine treatment led to nearly complete eradication of the tumor cells population (−97%±6%, *p*<0.0001).

We next generated PDAC experimental orthotopic tumors following engraftment of PC1.-0 cells in the pancreas of immune competent Syrian gold hamsters. Seven days later, tumors received an intratumoral injection of 900μg/kg of GMP CYL-02. Animals were treated at days 9, 11 and 13 following engraftment with 80mg/Kg gemcitabine, when control animals received NaCl 9^0/00^. Hamsters were euthanised15 days following tumor engraftment. *SSTR2, DCK* and *UMK* expression were monitored by qPCR in tumors as previously described^14^. Results shown Figure 5C demonstrates that CYL-02 transfection increases by 2 log the expression of the therapeutic genes into experimental PDAC tumors (*p*<0.005). We next performed immunohistochemistry (IHC) for SSTR2 expression, and found that approximatively 15 to 20% of tumor cells expressed the therapeutic transgene (Figure 5D). Last, we questioned the antitumoral activity of CYL-02 intratumoral injection, in the presence or not of systemic gemcitabine. Thus, experimentally growing tumors were injected with GMP CYL-02, then treated with gemcitabine as described before. Hamsters were euthanisedand tumor measured and sampled 8 days following gene transfer. We found that gemcitabine resulted in significant inhibition of tumor growth (−69%±10%, *p*<0.001), 8 animals out of 39 (20%) showing tumor regression (−25%±7%). CYL-02 by itself significantly inhibited tumor growth (−63%±13%, *p*<0.005), with 5 animals out of 36 (14%) showing tumor regression (−28%±9%). Remarkably, combining CYL-02 and gemcitabine showed strong inhibition of tumor growth as compared to control (−87%±10%, *p*<0.001), and superior antitumoral activity when compared to chemotherapy alone (−42%±8%, *p*<0.001). Combination therapy resulted in tumor regression (−31%±3%) in 31 out of 39 (79%) of the animals. We next analyzed the metastatic propensity of tumors from the different groups. We found that 86.5%±3% of animals from the control group presented with metastasis, mainly in the liver and in the peritoneum (Figure 5F). Gemcitabine or CYL-02 administration significantly reduced tumor dissemination (−41%±11% and 49%±27%, respectively, *p*<0.05). Here again, combination of CYL-02 and gemcitabine demonstrate strong inhibition of tumor dissemination (−81.5%±8%, *p*<0.005), and superior antimetastatic activity when compared to chemotherapy alone (−78%±6%, *p*<0.005). Collectively, these results demonstrate that CYL-02 significantly sensitizes tumor cells to chemotherapy, with more frequent tumor regression and less metastatic dissemination as compared to gemcitabine or CYL-02 administration alone. Last, we investigated the molecular mechanisms involved in the antitumoral activity of CYL-02 gene therapy when combined to gemcitabine chemotherapy. We found that tumor cell proliferation was significantly inhibited (PCNA labeling, −59±5%, *p*<0.05, Figure 5G), when cancer cell death by apoptosis was significantly increased (TUNEL assay, 11.25 ± 0.16-fold increase, *p*<0.001, Figure 5H). Collectively, we demonstrated herein that CYL-02 encoding for *SSTR2, DCK* and *UMK* cDNA complexed with PEI non-viral vector is safe and shows promising antitumoral and antimetastatic potential for the non-viral gene therapy of patients with PDAC.

## DISCUSSION

PDAC is characterized by a unique ability to withstand therapeutic aggression, and current treatments, mainly chemotherapies, are ineffective, as they increase survival times of patients only in weeks to months^2^. In previous studies, we demonstrated that *SSTR2, DCK* or *UMK* genes have strong potential to limit cancer cell proliferation, dissemination and to sensitize tumors to gemcitabine chemotherapy. This was achieved following *in vivo* intratumoral gene transfer using PEI non-viral vector in very aggressive PDAC experimental models^4,5,13^. These findings advocated for further clinical exploration, that would be preceded by the proper development and characterization of candidate gene therapy products.

During this process, we deliberately excluded viral vectors that may definitively improve therapeutic gene expression, but that may present with limitations, such as limited cloning capacity for adeno-associated virus, or even threats, illustrated by the risk of insertional mutagenesis due to host genome integration by lentiviral vectors, or supra physiological inflammation and risk of pancreatitis due to adenoviral transduction. As we found that long-term expression is not a priority for cancer therapy, and considering the unrivalled safety of non-viral vectors, we confirmed PEI as the gene delivery vehicle that would be used for the production of the gene therapy product later named CYL-02. Another technical innovation that would improve safety by limiting transgene expression to target cells is the use of original promoters with higher activity in tumor cells. Thus, each transgene within the plasmid DNA is under the control of hypoxia-sensitive promoters, namely glucose-responsive promoter (GRP) 78 (for *SSTR2* cDNA) and *GRP94* (for the DCK::UMK fusion cDNA), as PDAC cells are known to thrive for nutrients and to be frequently exposed to hypoxia^16^. In addition, these two promoters share common regulatory factors and are coordinately regulated into cells^17^, that is essential to avoid transcriptional competition. During this study, we generated a first generation of GTP in which the gene selection for bacterial selection could be expressed in eukaryotic cells. While this would not threaten further preclinical development, we solved this issue as this may jeopardize future clinical studies by modifying the GTP, then by performing “bridging” toxicology studies.

Safety has long been a primary concern in gene-therapy research, particularly after the death of a gene-therapy trial participant^18^. During this work, we identified that the MTD of CYL-02 in mice is of 1500 μg/kg of body weight. Pathology analysis further revealed mononuclear aggregates in the liver, that are generally found in mice and should not be attributed to the injection of a toxic substance, kidney congestion and bowel autolysis, probably as a result of agony before mouse euthanasia, and alveolar atelectasis, that usually corresponds to default in tissue fixing. This study was confirmed by the lack of toxicity of CYL-02 when injected in Syrian gold hamsters, including in tumors, as no animal death was recorded and control and CYL-02 injected animals grew with a similar trend. On the other hand, significant body weight loss, leukopenia and animal death were measured when CYL-02 was associated with gemcitabine. We found that these worrying signs were due to gemcitabine treatment, and that CYL-02 administration does not aggravate chemotherapy toxicity. We then explored in two different animal species whether CYL-02 may induce unwanted immune responses that could hamper translation to the clinic. We did not observe any evidence of acute nor systemic reaction following repeated injection of CYL-02, regardless of the route of administration. Collectively, we demonstrate that CYL-02 is safe, with a MTD of 1500μg/kg in mice and of 900μg/kg in hamsters when associated with chemotherapy, respectively. The safety factor generally applied to this type of molecule in oncology is 30, so to say that the calculated human dose equivalent (HED) for CYL-02 is 4 μg/kg of body weight. For a 62.5 kg male/female patient with PDAC, the minimum injected DNA dose to be tested would be 250μg of CYL-02. As one animal receiving 900μg/kg of CYL-02 showed frailty, we added an additional safety factor of 2, that converted into 2μg/kg in humans, leading to 125μg of the gene therapy product as the starting dose that was later administered to patients^14^. This dosage is consistent with early data from the use of the first generation CYL-02 in hamsters, 500μg/kg, where no adverse effects were observed after a single injection in tumor.

We next addressed the biodistribution of CYL-02 in mice and in preclinical models of PDAC. Importantly, we found that complexation with PEI does not allow for PCR amplification of the plasmid DNA (data not shown). This strongly suggests that positive signal by PCR corresponds to efficient delivery and release of the therapeutic DNA into cells. We demonstrate that CYL-02 is transiently detected in organs following i.v. injection, and is sequestered in the tumor following intratumoral injection with extremely low diffusion (<2.5% of the initial dose) in the vascular compartment. Importantly, CYL-02 was not detected in urine from hamsters following i.v. injection. This suggests that CYL-02 is not free, probably intraglobular, because it is not eliminated in the urine. This was later confirmed in humans receiving the GTP^14^. Interestingly, when used alone, CYL-02 is not detected in the lung, kidney and bowel. In association with gemcitabine, kidney and bowel show low levels of the gene therapy product, in sharp contrast with high levels of CYL-02 in spleen. We speculate that this may reflect microsomal release of the GTP caused by tumor cell lysis potentiated by gemcitabine, that is in line with the expected pharmacological action of the combination. Although toxicity studies showed CYL-02 to be safe, and although tissue biodistribution studies and respiratory system observations do not suggest any pulmonary toxicity, we nevertheless recommended pulmonary function monitoring in follow-up clinical studies, because the complex injected is considered as a nanovector, so that it can eventually forms micro-aggregates. Last, we recommend the monitoring of the cardiac, renal and hepatic functions, as we found traces of the GTP in these organs when injected at the highest dose.

In this work, we addressed for the first time the antitumoral potential of combining *SSTR2, DCK* and *UMK* gene transfer all together with chemotherapy treatment. We found that CYL-02 delivery resulted in comparable levels of expression of the three therapeutic genes, in cell lines and tumors, even if we identified a non-significant trend for lower expression of *DCK* and *UMK* as compared to *SSTR2*. This validate the choice of the two promoters that drive therapeutic gene expression. We found that 15 to 20% of cancer cells were transfected *in vivo* using CYL-02, that is in line with what we found using commercial PEI^4^. This validates that CYL-02 processing does not alter therapeutic gene delivery nor expression. Next, we demonstrated that CYL-02 treatment strongly sensitizes PDAC cells to chemotherapy, both *in vitro* and *in vivo* with increased antitumoral efficacy and more frequent tumor regression as compared to gemcitabine alone. Although comparisons are difficult to make, this surpasses the antitumoral effect following SSTR2 gene transfer^4^, or combination of *DCK::UMK* gene transfer and gemcitabine treatment^13^. In addition, combination therapy resulted in reduced capacity of tumor cells to disseminate, a property previously attributed to *SSTR2* expression^21^. The latter strongly suggests that the antitumoral properties of *SSTR2* and *DCK* and *UMK* are complementary to inhibit the growth of very aggressive PDAC tumors, and is preserved in the CYL-02 gene therapy product. Last, we found that CYL-02 combined with gemcitabine inhibits tumor cell proliferation and induces cell death by apoptosis. One of the major limitation of this work is that we didn’t use cutting-edge animal models of PDAC in this study, such as KPC mice^19,20^ as they were not available at the time this experiments were performed. These mice would also be instrumental to question the potential of CYL-02 and gemcitabine to induce an immune response against tumors.

Collectively, we provide during this work evidences that the CYL-02 gene therapy product generated under pre-GMP and GMP conditions for a pharmaceutical development is safe in preclinical models and strongly inhibits PDAC experimental growth when combined with gemcitabine. In addition, we determined the MTD that was later used to define the starting dose of CYL-02 to be used in humans^14^. In addition, this work provided with analytical procedures to explore the biodistribution of the GTP in patients, and also recommendation for organs monitoring during and after intratumoral gene delivery. Based on these findings, 57 patients with PDAC were treated by CYL-02 gene therapy combined with gemcitabine and no adverse events directly related to the action of the investigational drug were recorded. In some patients, CYL-02 and chemotherapy treatment resulted in tumor control, that advocated for a phase 2 study that was closed this year. Taken together, we demonstrate herein that rigorous and thorough investigations that are necessary to obtain authorization for clinical trial are instrumental and informative for the clinical transfer of gene therapy against PDAC.

## MATERIALS AND METHODS

### Animal studies

Experimental procedures performed on mice were approved by the ethical committee of INSERM CREFRE US006 animal facility and authorized by the French Ministry of Research: APAFIS#3600-2015121608386111v3. C57Bl/6 mice were obtained from Charles River and Syrian Gold hamsters from Harlan.

### Experimental tumor induction in Syrian gold hamsters

Hamster PDAC-derived PC-1.0 cells are grown in RPMI medium supplemented with 10% foetal calf serum, L-glutamine, antibiotics, antimycotics (Life Technologies), and Plasmocin^®^ (InvivoGen, Toulouse, France) in a humidified incubator at 37°C in 5% CO2. Six-week-old male Syrian gold hamsters were anesthetized by intraperitoneal injection of pentobarbital (80mg/kg) diluted in 0.9% NaCl, supplemented with oral anesthesia using oxygen/isofluorane (2.5 mixture) and PC-1.0 cells were implanted in the tail of pancreas as previously described^21^.

### Gene therapy product

CYL-02 is a complex of plasmid DNA and linear polymers of polyethyleneimine (in vivo-jetPEI® 22-kDa from Polyplus), prepared in 5% w/v glucose with a PEI nitrogen to DNA phosphate (N/P) ratio of 8 to 10. The plasmid within the gene therapy product encodes for DCK::UMK cDNAs (separated by the self-cleaving FMDV 2A peptide), and the human SSTR2 cDNA. Both DCK::UMK and SSTR2 cDNAs are under the coordinated transcriptional control of Glucose Responsive promoters (GRP78 and GRP94, respectively) that are highly sensitive to stress conditions, which prevails inside pancreatic tumors. The prokaryotic promoter-driven neomycin gene is used for bacterial selection and biodistribution and pharmacokinetic studies. The gene-therapy product is assembled and lyophilized by InvivoGen (Toulouse, France), following good medical product guidelines. Lyophilized CYL-02 was reconstituted by adding 2.5mL of sterile water for injectable preparation10 min before use. Therapeutic DNA and RNA were detected as described elsewhere^14^.

### *In vitro* analysis of CYL-02 activity and therapeutic efficacy

5×10e5 PC.1-0 cells were incubated with 1μg of therapeutic DNA and plated in 100-mm dishes. Three days later, culture medium was removed, cells were washed using PBS and cellular RNA was extracted using the RNAeasy kit from Qiagen. SSTR2, DCK and UMK expression was quantified by RTqPCR as described before^14^. For proliferation studies, 1×10e5 PC.1-0 cells were plated in 6-well plates in the presence of 1μg of CYL-02. Two days later, cells were treated with 20nM of gemcitabine and cells were counted 3 days later using the Z1 Coulter (Beckman).

### *In vivo* delivery of CYL-02 gene therapy product

Following laparotomy, tumors were measured using a caliper and randomized with mean=100mm^3^ in size as described before^21^. CYL-02 was administrated in in exponentially growing orthotopic tumors 8 days following tumor induction as previously described^21^. Control animals received 5% glucose. Gemcitabine (80 mg/kg) was injected i.p. every 2 days for 1 week, 48 hours after CYL-02 intratumoral injection. Control animals received 0.9% NaCl *i.p*. At 8 days post CYL-02 injection, animals were euthanized; primary tumors volume was measured using a caliper as described before^21^ and macrometastasis on lung, liver and peritoneum were counted.

### Experimental tumor analysis

PC.1-0 tumors were harvested and fixed in formalin. Four-micrometer-thick sections were prepared from paraffin-embedded sections and rehydrated. DNA fragmentation was performed using In situ Apoptosis Detection Kit according to the manufacturer’s indications (Takara Bio Inc.). For immunostaining, sections were incubated for 10 min in Protein Block, Serum-free reagent following antigen retrieval to reduce background staining (DakoCytomation). Slides were next incubated overnight at 4°C with anti-PCNA antibody (DakoCytomation, clone PC10, ref M0879, dilution: 1:100), or SSTR2 antibody (AbCAM clone [UMB1] ref ab134152, dilution: 1:100) diluted in Antibody diluent (DakoCytomation). Slides were washed and incubated in 3% H2O2 for 30 min at room temperature for endogenous peroxidase inhibition, quickly rinsed in distilled water, washed twice in PBS, and incubated for 30 minutes at room temperature with Envision+ system-HRP (DakoCytomation). After washing in distilled water, slides were incubated in AEC+ reagent and counterstained with Mayer’s hematoxylin. Immunostaining was recorded with an optical microscope, and quantified using VisioLab2000 image analyzer (Biocom). For each sample, fifteen fields were analyzed.

### Statistical analysis

Unpaired Student’s *t*-or Wilcoxon-Mann-Whitney tests were used to determine the statistical significance of differences between two groups using GraphPad Prism 9 software with the default settings. Methods of statistical analysis are indicated in the figure captions. Values are presented as \**p*□<□0.05, \*\**p*□<□0.01 and \*\*\**p*□<□0.005 and **** *p*□<□0.0001. Error bars are s.e.m. unless otherwise stated. The experiments were performed with a sample size *n* greater than or equal to three replicates. When monitoring tumor growth, the investigators were blinded to the group allocation but were aware of group allocation when assessing the outcome. No data were excluded from the analyses.

## Footnotes

1 NCT02806687

## ACKNOWLEDGMENTS

In loving memory of Prof. Gérard Tiraby (Invivogen) that was instrumental for this project. The authors thank Annie Souque for technical counseling and Severine Joubert for supervision counseling. Financial support: Région Midi-Pyrénées APRTCN 2006 N° 0401401 and APRTCN 2011 N° 12050667, ANR-RIB 07, Inserm Cossec and INVIVOGEN.

## AUTHOR CONTRIBUTIONS

Study concept and design: LB, PC, FG, HBT. Experimental procedures: OB, HL, NH, FV, GC. Data collection: LB, FG, PC, OB, HL. Data analysis: LB, FG, PC, OB, HBT. Supervision: LB, PC, FG, MT, SJ. Data interpretation: LB, FG, PC, HBT. Manuscript writing: PC. Manuscript revision: FG, LB.

## FIGURE LEGENDS

**Supplemental Figure 1.**
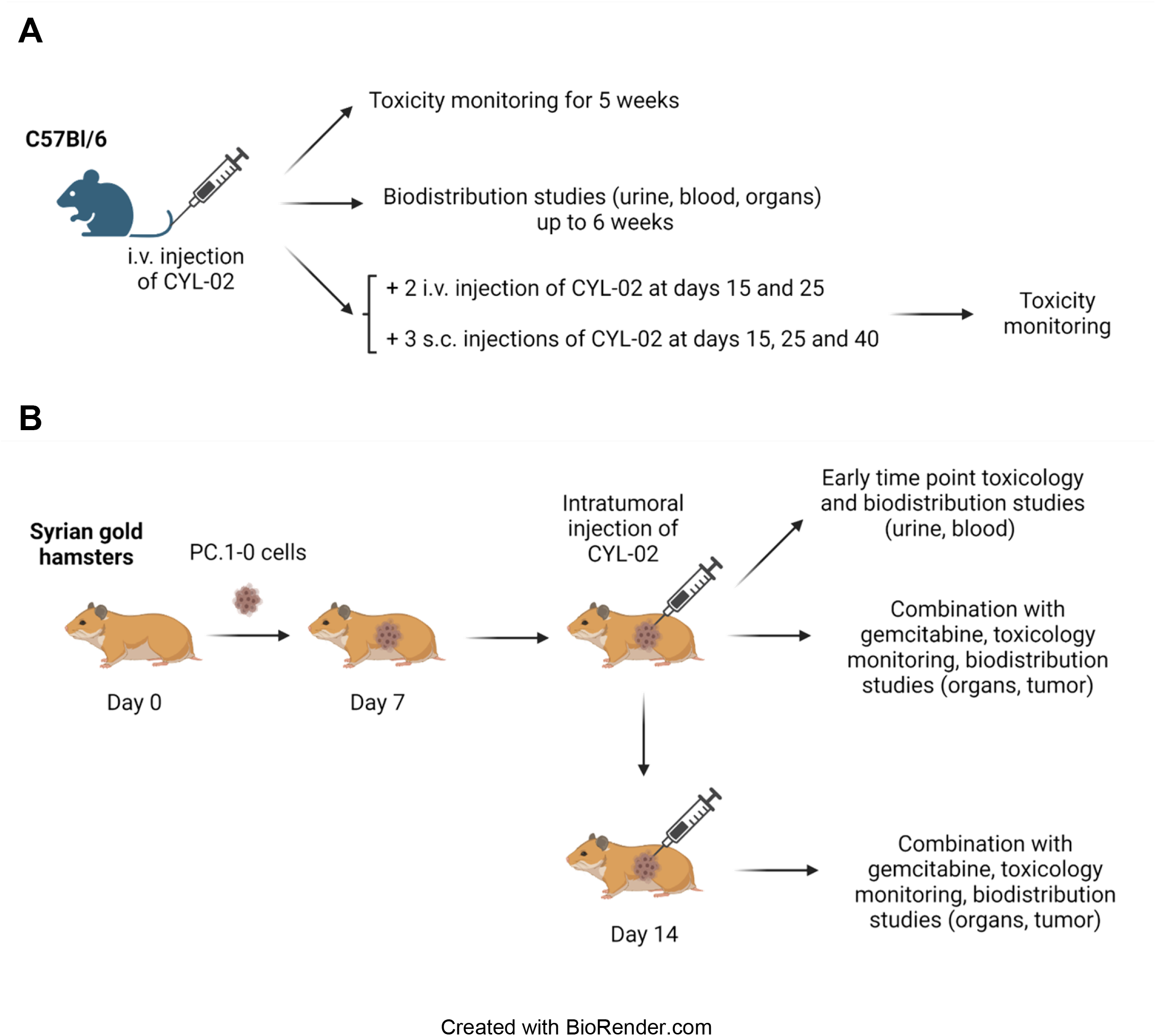
Schematic description of the preclinical animal models used during the study. Description of the toxicology and biodistribution studies following injection of CYL-02 in C57Bl/6 mice (A) and Syrian gold hamsters (B).

**Supplemental Figure 2.**
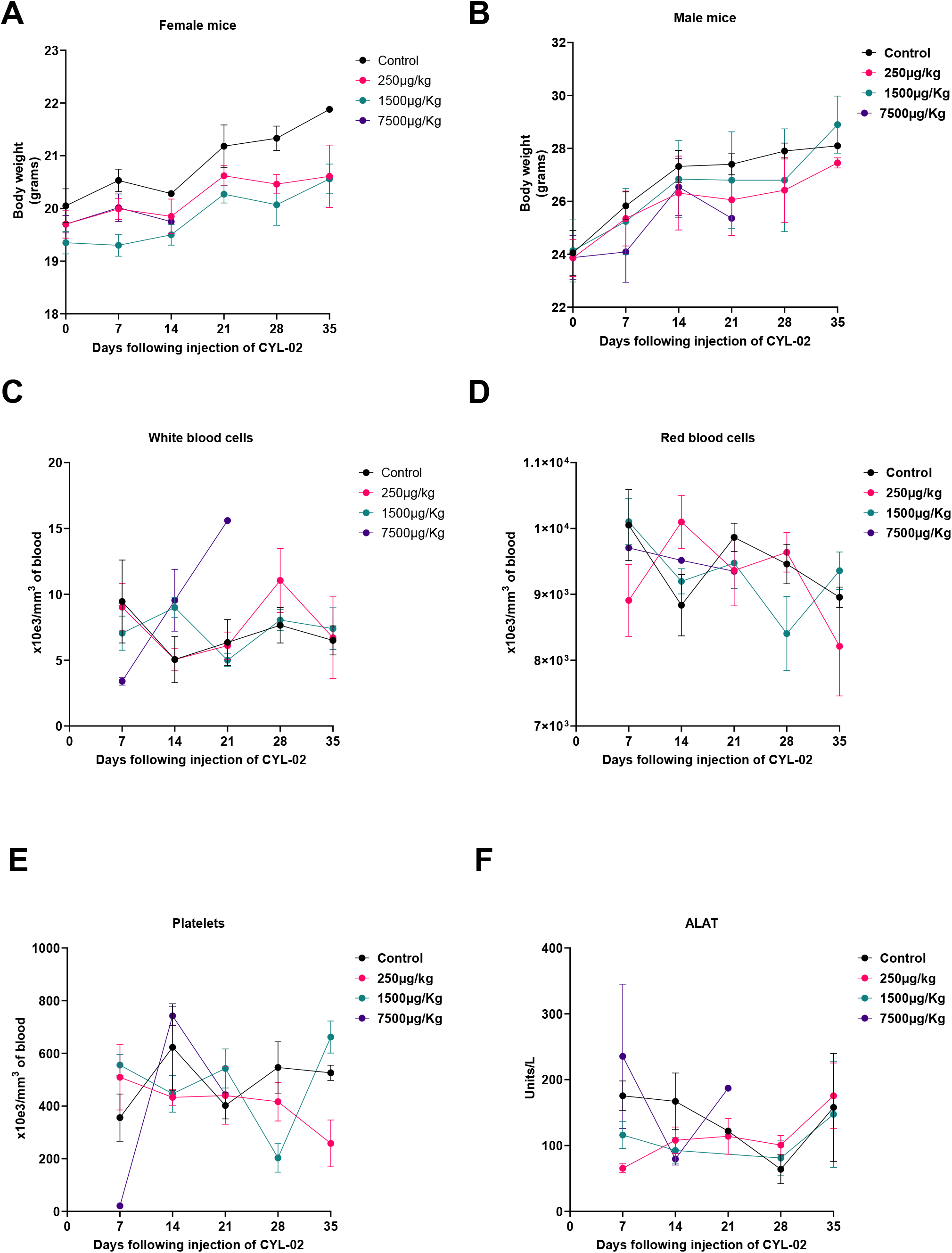
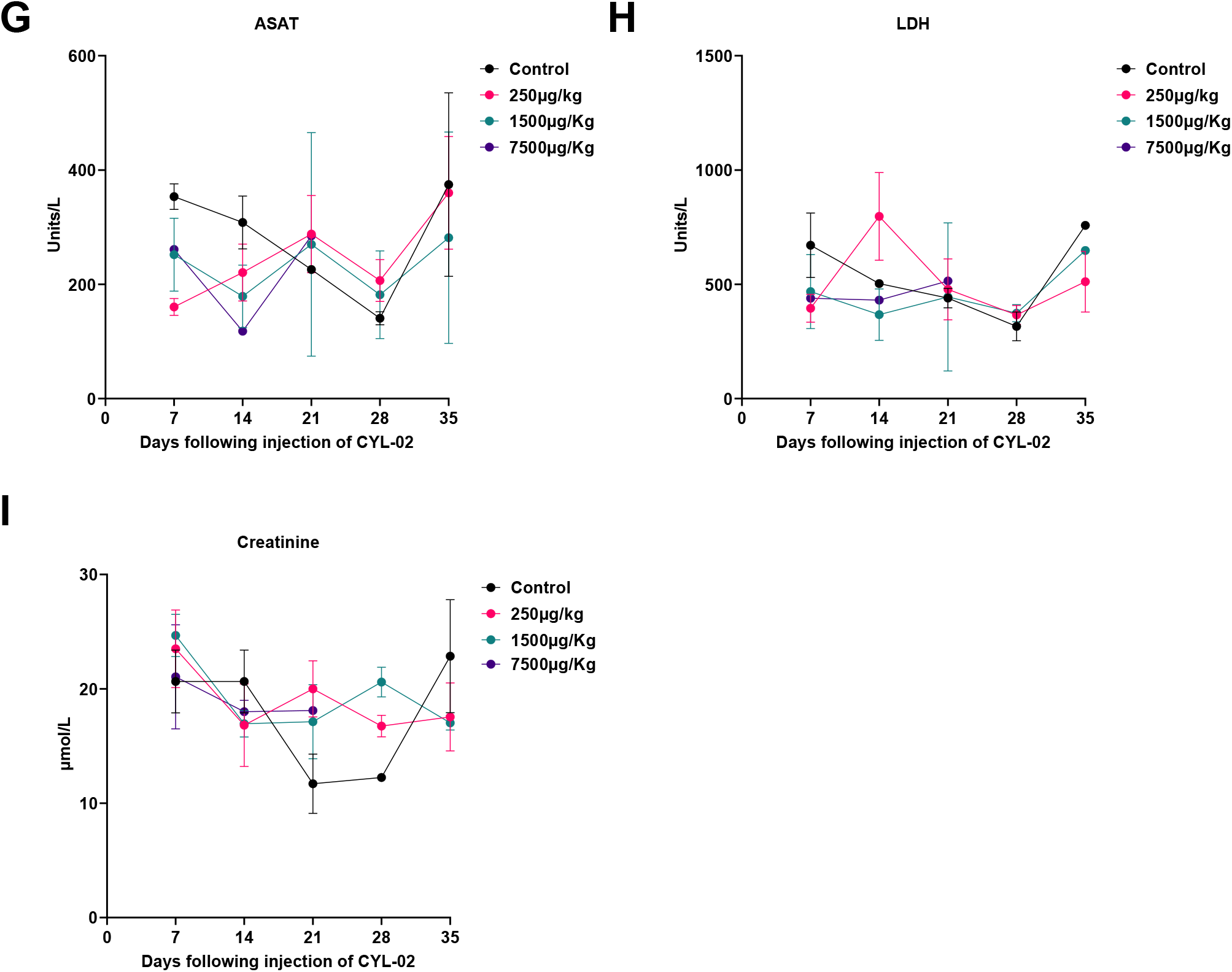
Toxicology monitoring in mice receiving an i.v. injection of CYL-02. 250μg/kg, 500μg/kg and 7500μg/kg of first generation CYL-02 was administered i.v. in C57Bl/6 mice from both sex. Body weight of female (**A**), or male mice (**B**), white (**C**), red (**D**) blood cells and platelets (**E**) counts, ALAT (**F**), ASAT (**G**), LDH (**H**) and creatinine (**I**) levels were monitored at the time indicated following injection of CYL-02. Results are mean ± SD of n=5 animals per group.

**Supplemental Figure 3.**
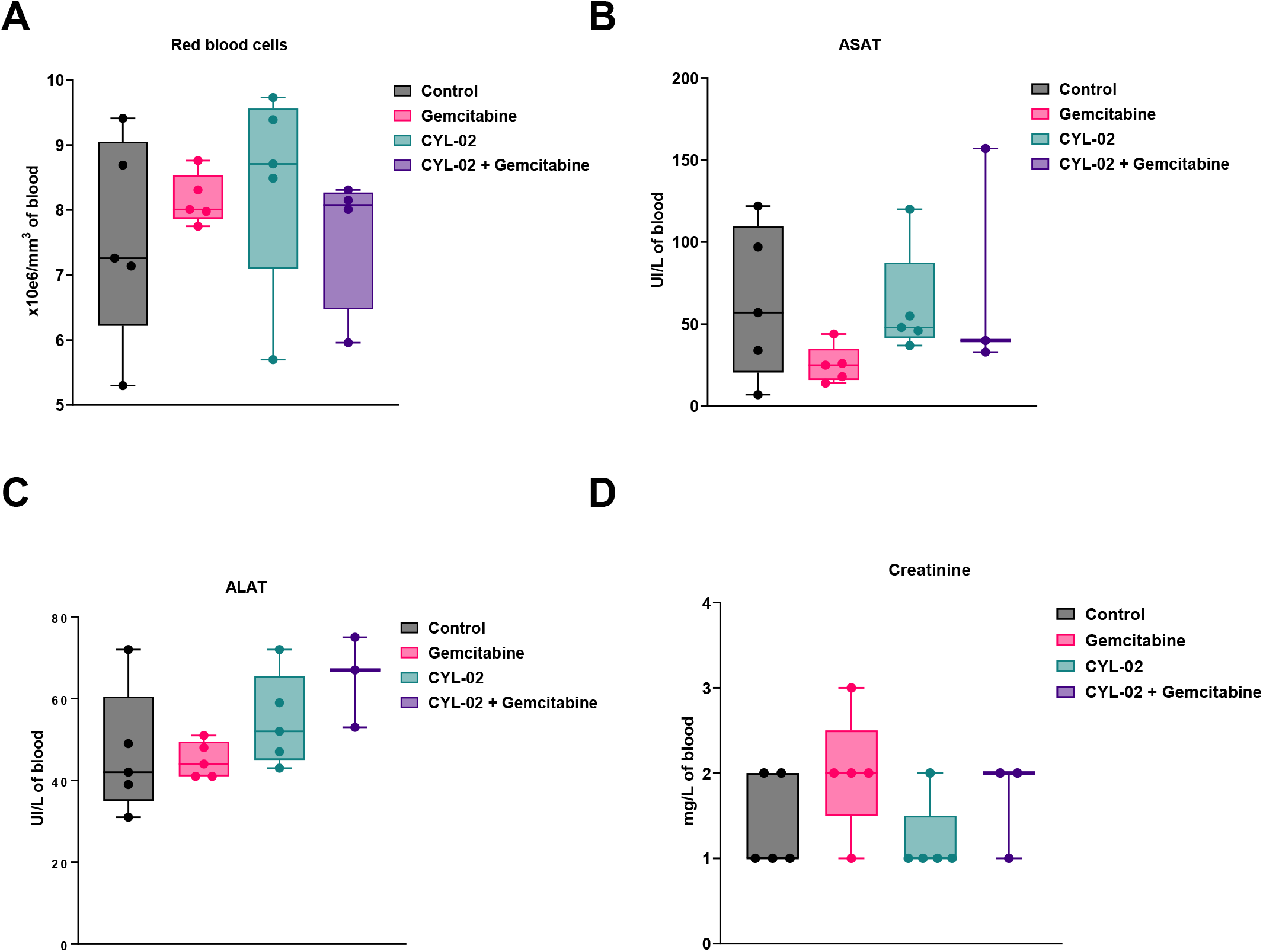
Toxicology monitoring in hamsters receiving an intratumoral injection of CYL-02 combined with gemcitabine treatment. Experimental orthotopic PDAC tumors were induced as described in Materials and Methods. Eight days later, 900μg/kg of second generation CYL-02 was administered in exponentially growing tumors, when control animals received 5% glucose. Gemcitabine was given at 80mg/kg i.p. every 2 days for a week. NaCl9^0/00^ was given i.p. as control. Red blood cells count (**A**), ASAT (**B**), ALAT (**C**), and creatinine (**D**) levels were monitored 7 days following injection of CYL-02 and treatment with gemcitabine. Results are mean ± SD of n=5 animals per group

**Supplemental Figure 4.**
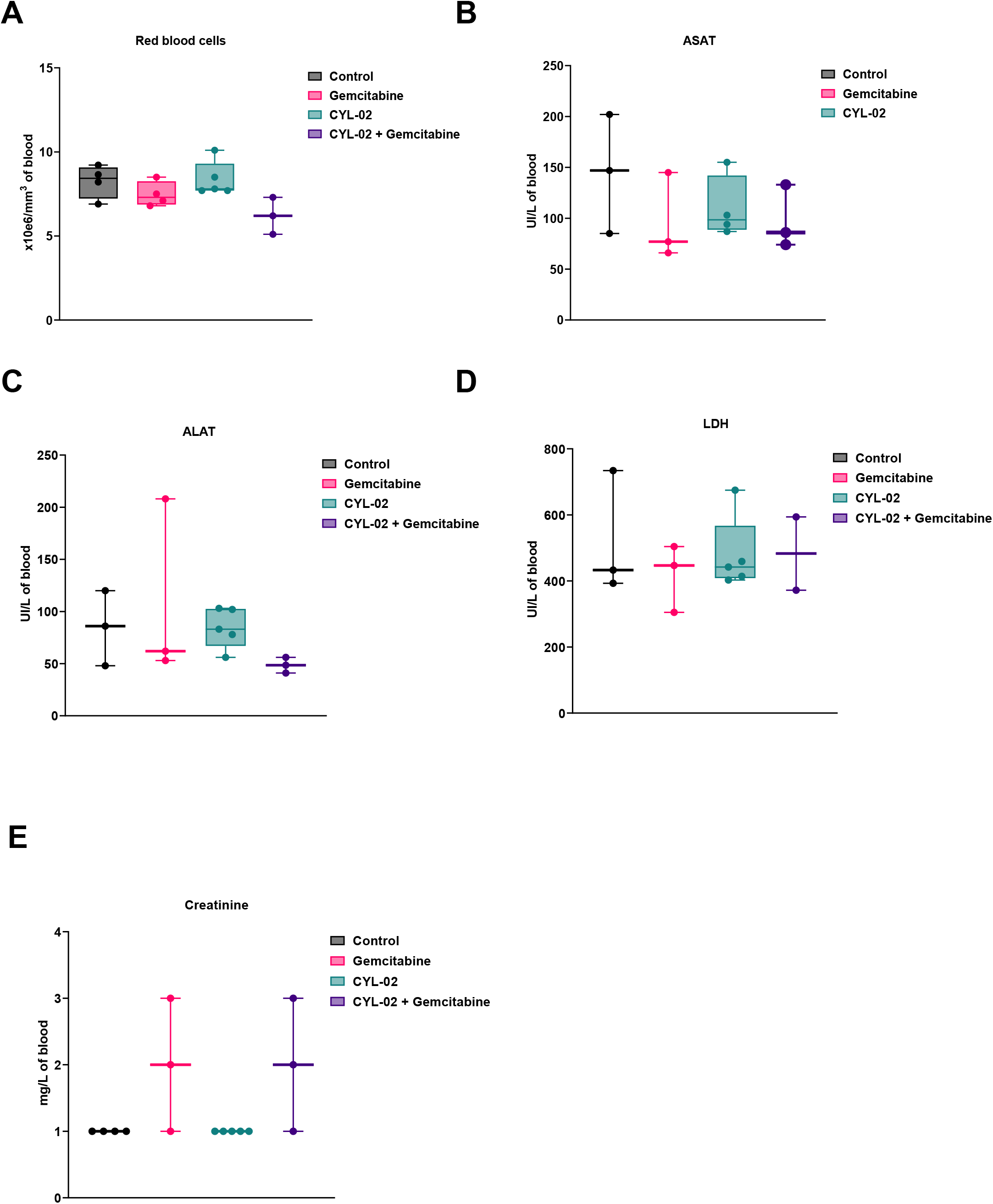
Toxicology monitoring in hamsters receiving two intratumoral injection of CYL-02 combined with gemcitabine treatment. Experimental orthotopic PDAC tumors were induced as described in Materials and Methods. 900μg/kg of second generation CYL-02 was administered at days 0 and 7 in exponentially growing tumors, when control animals received 5% glucose. Gemcitabine was given at 80mg/kg i.p. every 2 days for a week following each injection. NaCl9^0/00^ was given i.p. as control. Red blood cells count (**A**), ASAT (**B**), ALAT (**C**), LDH (**D**) and creatinine (**E**) levels were monitored 14 days following the first injection of CYL-02 combined with gemcitabine treatment. Results are mean ± SD of n=3 to 5 animals per group

## eTOC SYNOPSIS

